# Nutrient acquisition, rather than stress response over diel cycles, drives microbial transcription in a dessicated Namib Desert soil

**DOI:** 10.1101/432427

**Authors:** Carlos León-Sobrino, Jean-Baptiste Ramond, Gillian Maggs-Kölling, Don A Cowan

## Abstract

Hot desert surface soils are characterised by extremely low water activities for large parts of any annual cycle. It is widely assumed that microbial processes in such soils are very limited. Here we present the first metatranscriptomic survey of microbial community function in a low water activity hyperarid desert soil. Sequencing of total mRNA revealed a diverse and active community, dominated by *Actinobacteria*. Metatranscriptomic analysis of samples taken at different times over three days indicated that most functions did not fluctuate on a diel basis, except for a eukaryotic subpopulation which was induced during the cooler night hours. High levels of transcription of chemoautotrophic carbon fixation genes contrasted with limited expression of photosynthetic genes, indicating that chemoautotrophy is an important alternative to photosynthesis for carbon cycling in desiccated desert soils. Analysis of the transcriptional levels of key N-cycling genes provided strong evidence that soil nitrate was the dominant nitrogen input source. Transcriptional network analyses and taxon-resolved functional profiling suggested that nutrient acquisition processes, and not diurnal environmental variation, were the main drivers of community activity in hyperarid Namib Desert soil. While we also observed significant levels of expression of common stress response genes, these genes were not dominant hubs in the co-occurrence network.

## Background

Arid lands (deserts) are defined as having a level of precipitation (P) below the potential evapotranspiration (PET) level (P/PET < 1). Such lands cover an estimated one-third of Earth’s terrestrial surface (Laity, 2008) and are projected to expand in current climate change scenarios (Reich *et al.*, 2001). The Namib Desert, located along the western coast of Namibia and extending into southern Angola and northern South Africa, is the oldest (ca. 5 million years) continuously hyperarid (P/PET < 0.05) desert on Earth (Seely and Pallet, 2008).

According to current models, aridity results in habitat fragmentation, both geographically leading to “islands” of microbial biomass and diversity and temporally, producing long periods of functional inactivity (Pointing and Belnap, 2012; Collins *et al.*, 2014). However, recent evidence suggests that some activity is retained under these extreme conditions (Gunnigle *et al.*, 2017; Schulze-Makuch *et al.*, 2018). In recent years, the microbial ecology of various Namib Desert edaphic niches has been extensively studied (e.g. Scola *et al.*, 2017; Johnson *et al.*, 2017; Ronca *et al.*, 2015; Frossard *et al.*, 2015).

RNA sequencing has been employed to study microbial community functional patterns in many different aquatic and terrestrial ecosystems. The short life-span and high turnover of messenger RNA (Belasco and Brawerman, 1993) allows ephemeral states of microbial communities to be captured without significant interference from legacy biomolecules or inactive microbial populations, as might be the case in DNA- or protein-based studies (Nielsen *et al.*, 2006). In desert environments, active community changes over diel cycles or after rainfall have been described by 16S rRNA amplicon transcriptome studies (Gunnigle *et al.*, 2017; Štovíček *et al.*, 2017).

In this study, we analyzed 12 shotgun metatranscriptomes from hyperarid desert soils sampled over the course of 3 days. The experiment was designed to assess the diel transcriptional activity of desert edaphic microbial communities, particularly focusing on nutrient acquisition and stress response mechanisms. We also aimed to identify the key community members responsible for nutrient (C, N, P) cycling. Given the fluctuations in light, temperature and humidity to which desert soil communities are exposed within a daily cycle, we hypothesized that functional transcriptional profiles would also show distinct diurnal cycles.

## Results and Discussion

### Soil physicochemical characteristics

Soils were collected from a calcrete gravel plain site near the Gobabeb Research and Training Centre in the central Namib Desert (23°33′34′′S 15°02′25′′E) (Scholz, 1972) after a prolonged dry period (Supplementary Table S1). We implemented a three day sampling strategy with soil collection at near sunrise (6:00 h), at midday (12:00 h), at near sunset (18:00 h), and at midnight (24:00 h). Surface soil temperatures ranged from 21.4 ºC to 51.3 ºC, and soil air humidity ranged from 13% to 27.7% (Supplementary Fig. S2). An average photosynthetically active radiation (PAR) of 1722 ± 22 μmol photons m^−2^ s^−1^ was measured at 12:00 h, but was negligible or zero at 6:00, 18:00, and 24:00 h. Consistent soil respiration was recorded throughout the experiment (Supplementary Table S2).

The physicochemistry of soil samples was globally homogeneous. Localized heterogeneity was observed in four quadrats and related mostly to salt or phosphate concentration (Supplementary Table S2, Fig. S3).

### Library construction and sequence data

A sample from each time point was selected for library construction (n=12) based on homogenous soil physicochemical characteristics. Sequencing of cDNA libraries produced 285 million reads in total, with an average read length of 76 nt. After quality filtering and discarding rRNA and human-derived reads, 268 million high-quality reads were retained (Table S3).

### Taxonomic composition of the active soil microbial community

The taxonomic composition of the active microbial populations was similar throughout the study period (Supplementary Fig. S4), in spite of observed variations in temperature, humidity and light (Supplementary Fig. S2, Table S2). A single exception was the Day 2, 24:00 h library, which contained an unusually large proportion of fungal (19.5%, compared to an average of 2.8% in the remaining libraries) and *Firmicutes* (38.5%, compared to 5.9%) phylotypic sequences (Supplementary Fig. S4). This result most likely represented a random biological outlier, and this library was excluded from further analyses.

The phylogenetic analysis of the remaining 11 libraries showed that members of the domain Bacteria were most active (94.2 ± 2.6% of transcripts), with Eukarya comprising 4.3 ± 1.8% and Archaea 1.5 ± 1.0%. Virus-classified reads amounted to 0.04 ± 0.02% (Fig. 1A). Despite representing only a minor proportion, this is, to our knowledge, the first report of transcriptionally active viruses in dessicated hot desert soils, where lysogeny is considered to be the dominant state of virus populations (Zablocki *et al.*, 2015). However, the read volume was insufficient to provide a comprehensive survey of transcribed viral genes.

**Fig. 1:**
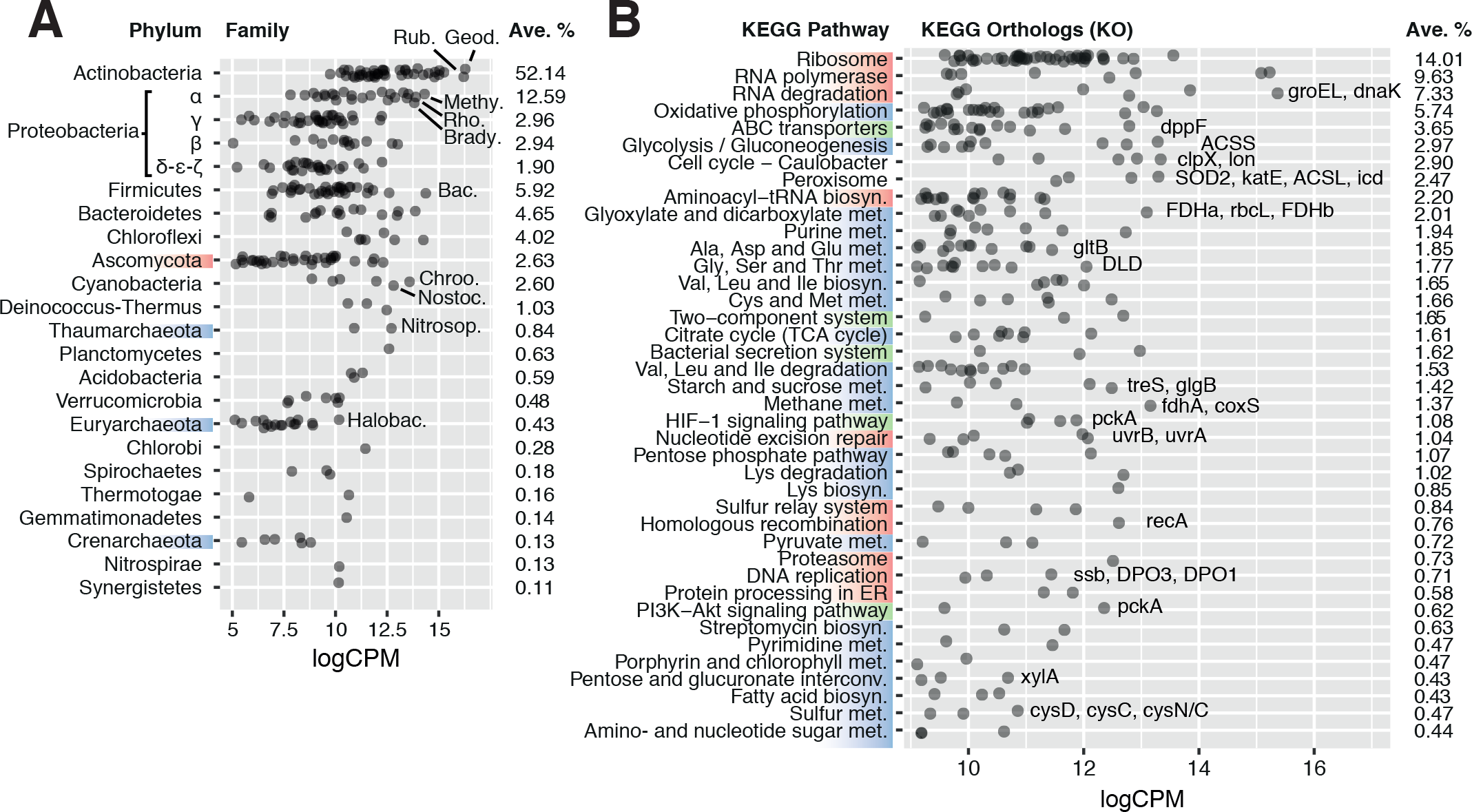
(A) Average transcriptional activity of the 20 most transcriptionally active microbial phyla and families. Phyla are sorted according to their average transcription levels (*aveLogCPM* function). Family transcript abundance is given in average log_2_ counts per million (logCPM). Geod.: *Geodermatophilaceae*; Rub.: *Rubrobacteriaceae*; Methy.: *Methylobacteriaceae*; Rho.: *Rhodobacteraceae*; Brady.: *Bradyrhizobiaceae*; Bac.: *Bacillaceae*; Chroo.: *Chroococcales*; Nostoc.: *Nostocaceae*; Nitrosop.: *Nitrosopumilaceae*; Halobac.: *Halobacteriaceae*. (B) Average transcript abundance of KEGG orthologs in the 40 most transcriptionally active KEGG pathways. Pathways are sorted according to their average log_2_ counts per million (*aveLogCPM* function). Upper KEGG classes are highlighted on the left axis by color: Genetic Information Processing (red), Metabolism (blue) and Environmental Information Processing (green). Categories Human Diseases and Organismal Systems were not included in the plot. Orthologs of particular interest are named, in order of transcript abundance, besides their respective pathway. Abbreviations: ACSL: acyl-CoA synthetase; ACSS: acetyl-CoA synthetase; clpX: Clp protease ATP-binding subunit; *coxS*: carbon-monoxide dehydrogenase small subunit; *cysD*: sulfate adenylyltransferase subunit 2; *cysC*: adenylylsulfate kinase; cysN/C: bifunctional enzyme CysN/CysC; DLD: dihydrolipoamide dehydrogenase; *dnaK*: molecular chaperone DnaK; DPO1/3: DNA polymerase I/III; *dppF*: dipeptide transport system ATP-binding protein; FDHa/b: formate dehydrogenase alpha/beta subunit; *fdhA*: formaldehyde dehydrogenase; *glgB*: 1,4-alpha-glucan branching enzyme; *gltB*: glutamate synthase; *groEL*: chaperonin GroEL; *icd*: isocitrate dehydrogenase; *katE*: catalase; *lon*: Lon protease; *pckA*: phosphoenolpyruvate carboxykinase; *rbcL*: ribulose-bisphosphate carboxylase large chain; *recA*: recombination protein RecA; SOD2: superoxide dismutase; *ssb*: single-strand DNA-binding protein; *treS*: maltose alpha-D-glucosyltransferase / alpha-amylase; *uvrA*/*B*: excinuclease ABC; *xylA*: xylose isomerase.

Seven bacterial phyla (*Actinobacteria*, *Proteobacteria*, *Firmicutes*, *Bacteroidetes*, *Chloroflexi*, *Cyanobacteria*, and *Deinococcus-Thermus*) and one eukaryal phylum (*Ascomycota*) each contributed more than 1% of the classified reads, jointly comprising 93.1 ± 2.3% of the total active community (Fig. 1A and Supplementary Fig. S4). The most active phylum globally was Actinobacteria, producing 52.1 ± 5.4% of the classified transcripts, followed by Proteobacteria, with 20.2 ± 2.6% of the transcripts. Two prominent actinobacterial families, *Geodermatophilaceae* and *Rubrobacteraceae*, represented 8.2 ± 1.8% and 7.0 ± 1.9%, respectively (Fig. 1A). Both families have been routinely detected in desert soils and their members typically exhibit high stress tolerance and are metabolically versatile (Rainey *et al.*, 2005; Favet *et al.*, 2013; Sghaier *et al.*, 2016; Albuquerque and da Costa, 2014; Normand *et al.*, 2015).

### Functional profile of the microbial community

All core metabolic pathways were transcribed (Fig. 1B), including replication genes, indicating that the active fraction of the soil microbial community had complete functionality.

This observation implies the existence of a xeroresistant microbial community in this hyperarid desert niche. Tolerance; i.e., survival with impaired or no activity and no growth, is regarded as the most common strategy adopted by microbial communities under extreme xeric stress, such as in hyperarid desert soils (Lebre *et al.*, 2017). Hyperaridity results in habitat fragmentation and concentrates activity in sheltered “islands of fertility” and during brief wet periods (Pointing and Belnap, 2012; Collins *et al.*, 2014). Active microbial populations have recently been detected in hyperarid soils from the Atacama Desert (Schulze-Makuch *et al.*, 2018), although at very reduced activity levels that suggest a temporally or metabolically limited state. However, our transcription results, which demonstrate the presence of a functional fraction of the microbial community, suggest that resistance, rather than tolerance, is a strategy adopted by some of the resident taxa. Resistance is here defined as the maintenance of function, despite the impositions of extreme environmental parameters (i.e., hyperaridity) (Harrison *et al.*, 2007).

The coexistence of dessication-resistant and -tolerant microbial taxa has been observed in non-arid soils, where Actinobacteria remain active during dry periods whereas Acidobacteria become the dominant active group upon rewetting (Barnard *et al.*, 2013). Our findings therefore extend this dual-response model to soils under extreme xeric stress.

Genes encoding elements of stress resistance and damage repair mechanisms were highly transcribed. Chaperone genes *groEL* and *dnaK* (4.9% and 1.5% of the classified transcripts, respectively) (Fig. 1B), and protease genes involved in protein quality control (e.g., *clpX/P* and *lon*; 1.7% and 0.7%, respectively) were among the most transcribed. Furthermore, the high relative abundances of peroxisomal orthologs (2.4%), such as superoxide dismutase (SOD) and catalase (*katE*), as well as DNA repair gene transcripts (*recA*, *uvr*, Fig. 1B), support the widely held view that radiation- and desiccation-induced damage (particularly related to oxidation processes) are the major stresses for microbial cells in hyperarid hot desert soils (Makhalanyane *et al.*, 2015). The production of compatible solutes and capsule formation, which are common microbial adaptation mechanisms for desiccation tolerance (Lebre *et al.*, 2017), were suggested by polysaccharide and trehalose biosynthesis gene transcripts such as the alpha-glucan branching enzyme gene *glgB* (0.4% of transcripts) (Rashid *et al.*, 2016) (Fig. 1B). Overall, the transcriptional profile of the microbial community coherently reflects known strategies of desiccation resistance predicted from genomic analyses of desiccation-tolerant microorganisms (Lebre *et al.*, 2017; Schulze-Makuch *et al.*, 2018), allowing full activity of a subpopulation throughout hyperarid periods.

### Nutrient cycling and key active taxa

Carbon, nitrogen, and phosphorus are the major limiting nutrients for microbial communities, and for oligotrophic desert soil communities in particular (Cleveland and Liptzin, 2007; Delgado-Baquerizo *et al.*, 2013; Johnson *et al.*, 2017). The transcriptional activity of orthologs involved in assimilation pathways of these nutrients was thus investigated, to evaluate the contributions of specific taxa to these processes in the community.

High levels of functional redundancy were evident (Fig. 2). However, some important ecosystem functions appeared to be taxon-specific. For example, nitrate reductase (*nar*) genes, which encode key enzymes in nitrogen assimilation in soils (Merrick and Edwards, 1995) (Fig. 3A), were transcribed almost exclusively by members of the *Nitrospiraceae* family (Fig. 2A), indicating that this family plays a key role in the nitrogen cycling of Namib desert soil communities.

**Fig. 2:**
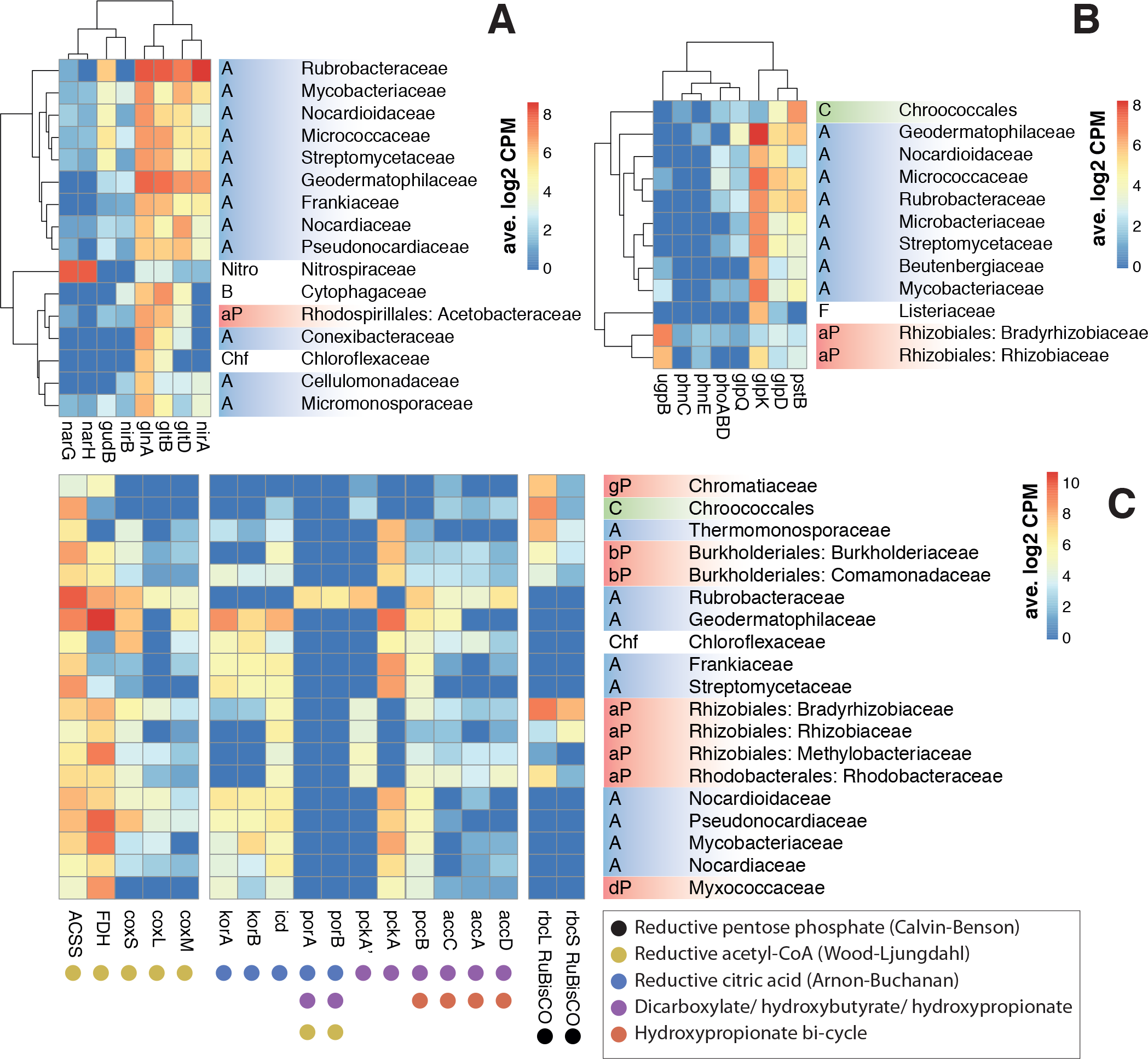
Correlation between family-level taxonomy and key KEGG ortholog transcription from nitrogen (A), phosphorus (B) and carbon (C) assimilation pathways in selected prokaryotic taxa. (A) Nitrogen metabolism orthologs: *gln*: glutamine synthetase; *glt*: glutamate synthase; *gud*: glutamate dehydrogenase; *nar*: nitrate reductase; *nir*: nitrite reductase. (B) Phosphorus assimilation and G3P metabolism orthologs: *glpD*: glycerol 3-phosphate dehydrogenase; *glpK*: glycerol kinase; *glpQ*: glycerophosphodiester phosphodiesterase; *phn*: phosphonate transport; *phoABD*: alkaline phosphatase; *pst*: phosphate transport; *ugp*: sn-glycerol-3-phosphate transport. (C) Carboxylase gene orthologs involved in carbon fixation pathways: ACSS: acetyl-CoA synthase; *acc*: acetyl-CoA carboxylase; FDH: formate dehydrogenase; *icd*: isocitrate dehydrogenase; *kor*: 2-oxoglutarate synthase; *cox*: CO dehydrogenase; *pcc*: acetyl/propionyl-CoA carboxylase; *pck*: PEP carboxylase; *por*: pyruvate synthase; *rbc*: RuBisCO. Phylum abbreviations: A : *Actinobacteria*; B: *Bacteroidetes*; C: *Cyanobacteria*; Chf: *Chloroflexi*; F: *Firmicutes*; Nitro: *Nitrospirae*; P: *Proteobacteria* (a: alpha, b: beta, d: delta, g: gamma). Only families with > 6 average log_2_CPM for the selected genes were included in A and B. Hierarchical clustering of rows and columns was performed with *hclust* function.

**Fig. 3:**
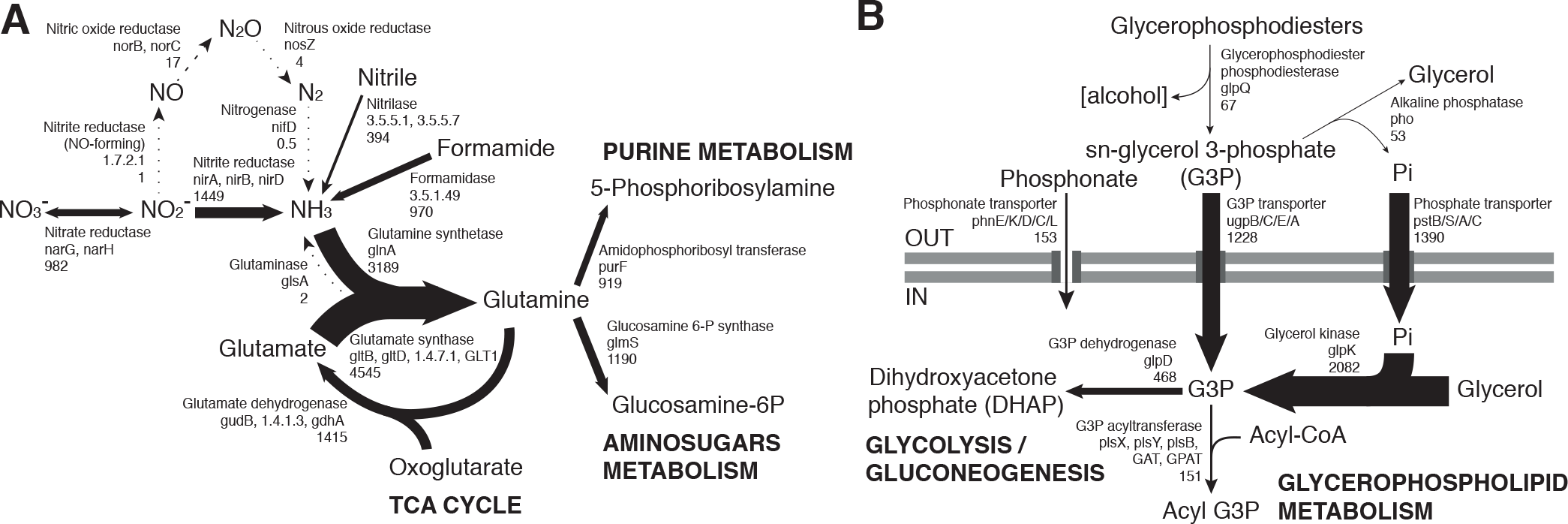
Community-level transcription of nitrogen (A) and phosphorus (B) assimilation pathways, including glycerol phosphate metabolism. Gene codes in bold highlight the most abundant orthologs from a group performing the same function, when there is a large transcript abundance difference. Numbers below gene codes show the total average counts per million (cpm) of all orthologs. Arrow thickness in each figure is proportional to the indicated cpm value.

### Nitrogen assimilation

Nitrogen-fixing bacterial taxa such as *Geodermatophilaceae*, *Frankiaceae* and *Rhizobiales* (Merrick and Edwards, 1995; Sellstedt and Richau, 2013) were among the most active taxa (Fig. 1A). However, transcripts relating to the nitrogen metabolism KEGG pathway represented a small portion (0.2%) of our soil metatranscriptomes and virtually no *nifD* nitrogenase transcripts were detected (Fig. 3A). These findings are compatible with recent observations that hypolithic communities, and not surface soil communities, were the primary sources of N_2_-fixation in Namib Desert gravel plains (Ramond *et al.*, 2018).

Transcriptome data suggested that nitrate reduction, most transcribed by the *Nitrospiraceae* family, and nitrite reduction primarily transcribed in actinobacterial taxa (*nar* and *nir* genes, respectively, Fig. 2A) were the dominant processes in the generation of biologically available nitrogen in the community from a NO_3_^−^ and NO_2_^−^ reservoir (Fig. 3A). These nitrogen species may be accumulated in soils during infrequent wet periods, possibly as a result of the activation of genes and microorganisms inhibited during dessicated conditions (Scherer *et al.*, 1984), or from dry atmospheric deposition processes (Báez *et al.*, 2007; Jia *et al.*, 2016).

### Phosphorus and Sulfur assimilation

Most phosphorus is available to soil microbial communities as inorganic phosphate (Pi), solubilized from the mineral soil fraction or released from organic molecules by the alkaline phosphatase (White and Metcalf, 2007). *Pst* phosphate transporter gene transcripts were abundant in the community (1390 average counts per million, cpm). Organic phosphate sources were also possibly exploited, as suggested by transcription of the *phn* phosphonate transporter gene (153 cpm) and especially the sn-glycerol 3-phosphate (G3P) transporter gene *ugp* (1228 cpm) (Fig. 3B). Although the expression of *phn* and *ugp* can be inhibited by Pi (Schowanek and Verstraete, 1990; Brzoska *et al.*, 1994), organic P utilization may still be an important microbial community trait in oligotrophic desert environments (Vikram *et al.*, 2016). The *ugp* genes were principally transcribed by members of the Order *Rhizobiales* (Class *alpha-Proteobacteria*), potentially replacing Pi transport as a phosphorus acquisition mechanism (Fig. 2B). Plant exudates or membrane phospholipids are possible sources of G3P in soils (Collins *et al.*, 2014; Lidbury *et al.*, 2017; Ding *et al.*, 2012). The *glpQ* gene product can cleave these compounds, releasing G3P and triggering activation of the *ugp* transporter genes (Brzoska and Boos, 1988).

*GlpQ* is an extracellular enzyme which has been implicated in cooperative interactions between proteobacteria (Lidbury *et al.*, 2017). In our dataset, *glpQ* was mostly transcribed in *Actinobacteria*. The most active actinobacterial family *Geodermatophilaceae*, however, transcribed *glpQ*, but not the G3P transporter *glp* or the alkaline phosphatase *pho* genes, which would dephosphorylate G3P, releasing Pi for its own consumption (Fig. 2B). Our data therefore suggest a putative interaction between *Geodermatophilaceae* and *Rhizobiales*, with the former providing access to phosphorus as G3P for the latter. This interaction may be of considerable importance for desert community maintenance, as both taxa were amongst the most transcriptionally active (8.2% and 6.1%, respectively) (Fig. 1A).

Reductive sulfate assimilation (*cys* genes) was the dominant transcribed S-cycling pathway in the community. However, transcripts for the sulfate transporter *cysPUWA*, were mostly associated to proteobacteria, particularly the Burkholderiales family, suggesting a central role of this group in sulfur assimilation and cycling.

### Carbon fixation

A defining feature of arid soils is low productivity and a low organic carbon content (Delgado-Baquerizo *et al.*, 2013). Hyperaridity imposes severe constraints on oxygenic photosynthesis, for which water is the electron donor (Warren-Rhodes *et al.*, 2006). Furthermore, soil communities outside of sheltered fertile islands (i.e., hypoliths, endoliths, or biological soil crusts) typically have a very low abundance of the phototrophic cyanobacteria (Stomeo *et al.*, 2013; Makhalanyane *et al.*, 2013). Perhaps not surprisingly, transcription of photosynthetic pathway genes and phototrophic organisms was limited in our dataset (Fig. 1B, Fig. 1A). Notably, reads classified within the Glyoxylate and Dicarboxylate pathway (2.0%) exceeded those assigned to photosynthetic KEGG pathways and, surprisingly, also significantly exceeded those from the TCA Cycle (0.3% and 1.6%, respectively, two-tailed t-test p < 0.005) (Fig. 1B). We also observed a higher number of transcripts assigned to acetyl-CoA synthetase (ACSS, 9874 cpm) and formate dehydrogenase (FDH, 10254 cpm) compared to RuBisCO (*rbcL*/*S*, 2672 cpm) (Fig. 1B). These observations strongly suggest that chemoautotrophic (“dark”) carbon fixation and/or CO_2_ reassimilation mechanisms are important microbial processes by which inorganic C enters the soil microbial community. Chemoautotrophic carbon fixation has been shown as an important process in marine environments, even when photosynthesis is active (Palovaara *et al.*, 2014; Aylward *et al.*, 2015), and is a potentially major process in soils (King and Weber, 2007; Pratscher *et al.*, 2011). Therefore, we examined the activity of carboxylase genes in greater depth, and their distribution between different families of microorganisms.

Although RuBisCO gene transcripts (*rbc*) from the Calvin-Benson-Bassham (CBB) cycle were significant (average 2672 cpm) (Fig. 1B), the majority were assigned to non-photosynthesizing *alpha-Proteobacteria* rather than to *Cyanobacteria* (Fig. 2C). This suggested that the CBB cycle acted predominantly in chemoautotrophic CO_2_ fixation or as an electron sink (Badger and Bek, 2008; McKinlay and Harwood, 2010), rather than in photosynthesis.

Orthologs of the acetyl-CoA synthase ACSS (9874 cpm), CO dehydrogenase *coxS* (1701 cpm) and formate dehydrogenase FDH (10254 cpm) genes, involved in the reductive acetyl-CoA cycle (Wood-Ljungdahl pathway), were significantly transcribed in a wide range of taxa (Fig. 2C). These carboxylases are widely distributed in soil bacteria (King and Weber, 2007) and are active in desert actinobacteria (Sghaier *et al.*, 2016). The activity of these genes in Namib Desert soil microbial communities may be related to the very low energy requirements and the capacity to coassimilate one-carbon compounds or acetate of this pathway (Fuchs, 2011), making it well suited to oligotrophic niches.

The reductive citric acid cycle (Arnon-Buchanan cycle) carboxylases *kor* (2-oxoglutarate synthase) and *icd* (isocitrate dehydrogenase) were transcribed by many actinobacterial families, but surprisingly not by *Rubrobacteraceae* (Fig. 2C). Instead, *Rubrobacteraceae* transcribed the acetyl and propionyl-CoA carboxylase *pccB* and *accA/C/D*, the phosphoenolpyruvate (PEP) carboxylase *pckA* and the pyruvate synthase *por*, which participate in other pathways of chemoautotrophic carbon fixation (Fuchs, 2011) (Fig. 2C). These pathways could only be partialy detected in *Rubrobacteraceae*, as several key genes (e.g., malonyl-CoA reductase, 4-hydroxybutyryl-CoA dehydratase) were not identified. These pathways, either full or partial, allow prokaryotes to coassimilate reduced and uncommon C compounds and to fix carbonate (Fuchs, 2011; Zarzycki and Fuchs, 2011). Our results therefore suggest that the actinobacterial *Rubrobacteraceae* family may be important in inorganic carbon acquisition in desert soils, partly due to a high plasticity in chemoautotrophic metabolism.

### Circadian differential gene expression

Environmental variations often cause microbial communities to exhibit differential activity profiles over temporal timescales (e.g., daily or seasonally), both in phototrophic and non-phototrophic groups (Klatt *et al.*, 2013; van der Meer *et al.*, 2005; Ottesen *et al.*, 2013, 2014; Aylward *et al.*, 2015).

Diel transcriptional periodicity was examined using EdgeR (Robinson *et al.*, 2010). Time pairs were contrasted independently, as well as “day” (12:00 and 18:00) versus “night” (24:00 and 6:00) groups as defined by their contrasted temperature and humidity records (Supplementary Fig. S2). Interestingly, pairwise comparisons identified no differentially expressed orthologs (p > 0.05) between the 12:00 and 18:00 or between the 24:00 and 6:00 datasets. When “day” and “night” data were contrasted, 13 of 2265 orthologs (0.57%) were significantly (p <0.05) induced during the night (Supplementary Table S4). None were highly transcribed orthologs, suggesting that under extreme dry conditions, desert soil communities are generally functionally stable and that their principal functions are not regulated on a diel scale. This conclusion has implications in terms of the perceived drivers of microbial community function, as our results suggest that the constant xeric stress is a more significant driver of *in situ* functionality than daily environmental variations (temperature, air moisture or light). We predict that this functional stability would only be substantially disrupted by stochastic events such as rainfall, which is recognised as a main driver of community assembly and activity in arid soil environments (Belnap *et al.*, 2005; Pointing and Belnap, 2012; Frossard *et al.*, 2015; Scola *et al.*, 2017).

Surprisingly, we observed a marked enrichment in differentially transcribed eukaryal orthologs, including tubulin, dynein, myosin, SF3B, *dnaJ* and ANP1 genes (Supplementary Table S4). This suggested that the active fungi, which only represented 2.8% of the total transcripts, were most active during the cooler and higher atmospheric humidity night hours, contrary to the generally stable activity pattern observed for the rest of the community. This is consistent with observations made on fungi and lichens from arid environments, which appear to grow optimally during small air moisture pulses (Palmer *et al.*, 1987; Jacobson *et al.*, 2015).

### Transcriptional network analysis

The temporal co-variation of KOs was determined in order to examine whether coordinated patterns of gene transcription existed within the soil community. 624 orthologs were used to construct a transcriptional network, 83.3% of which (520) clustered into 4 distinct modules (A to D, Fig. 4). The larger clusters A and B were composed of positively interrelated orthologs, although no specific functional enrichment within each module was observed.

**Fig. 4:**
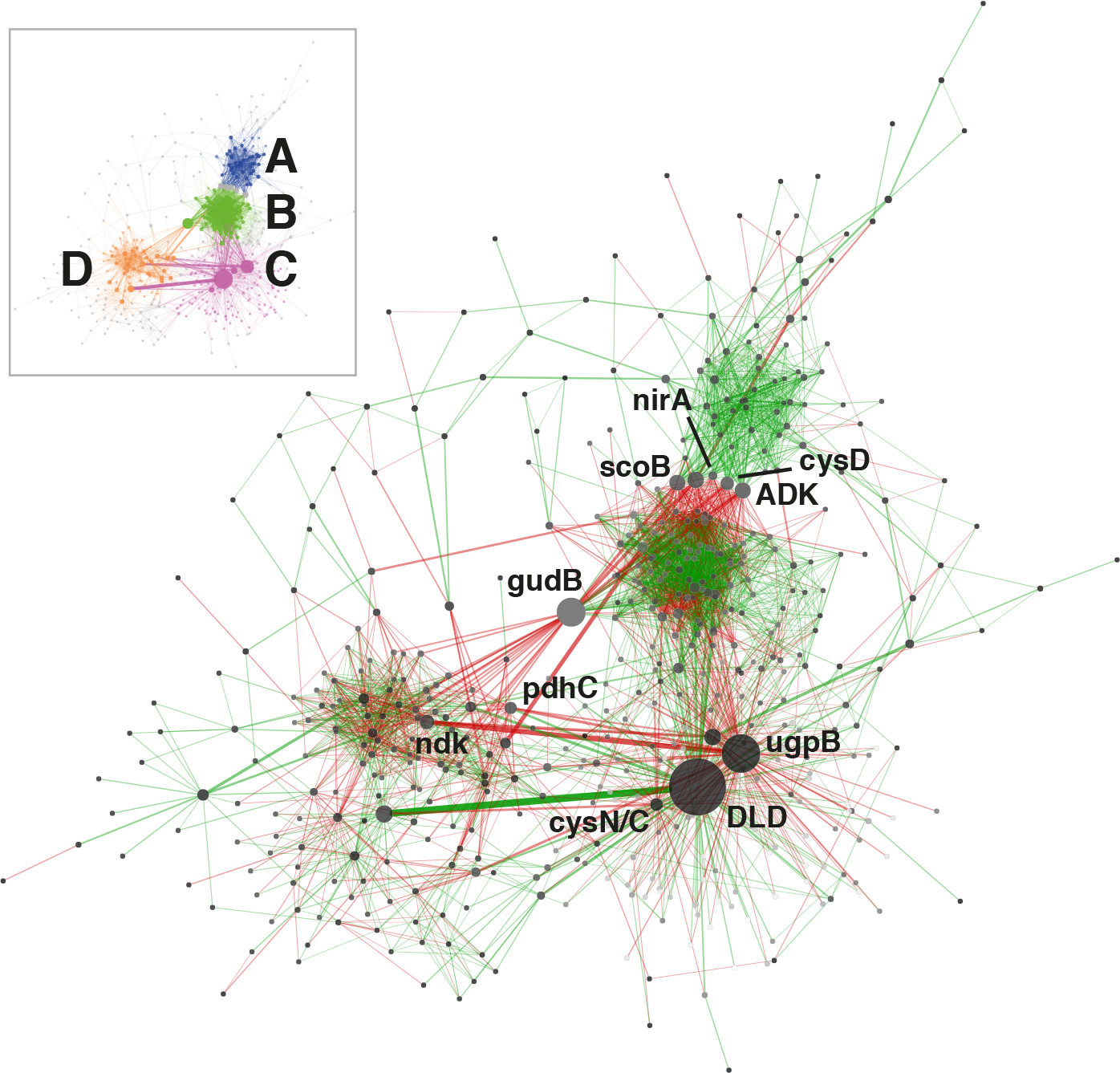
KEGG ortholog transcriptional network. Only orthologs with average abundances of log_2_ cpm > 7 in the complete dataset were used for computation. Node size is proportional to betweenness centrality and edge thickness is proportional to betweenness. Green or red edge lines indicate a shared positive or negative correlation, respectively. Ortholog abbreviations: ADK: adenosine kinase; *cysD*: sulfate adenylyltransferase (sulfate-activating complex); *cysN/C*: bifunctional enzyme *CysN/CysC* (sulfate-activating complex); DLD: dihydrolipoamide dehydrogenase; *gudB*: glutamate dehydrogenase; *ndk*: nucleoside-diphosphate kinase; *nirA*: ferredoxin-nitrite reductase; *pdhC*: pyruvate dehydrogenase; scoB: 3-oxoacid CoA-transferase; *ugpB*: sn-glycerol 3-phosphate transport system.

The principal transcriptional network clusters were connected through three orthologs: the dihydrolipoamide dehydrogenase DLD, the G3P transporter subunit *ugpB* and the glutamate dehydrogenase gene *gudB* genes. These genes occupy network hub positions and therefore changes in their transcriptional status could result in large shifts in community function. The dominant metabolic function associated with the highly transcribed (3917 cpm, Fig. 1B) DLD gene is in the TCA cycle, but has also been shown to affect sugar transport and capsule formation via direct interactions with membrane transporters (Tyx *et al.*, 2011). The importance of exopolysaccharides in desiccation resistance (Lebre *et al.*, 2017), and the TCA cycle in carbon metabolism regulation, support the centrality of DLD in the community network. The gene *ugpB*, as previously discussed, was linked to the rhizobial community as part of a nearly exclusive phosphorus assimilation mechanism (Fig. 2B). The *gudB* gene product catalyzes the synthesis of glutamate, the principal acceptor metabolite in NH_3_ assimilation (Merrick and Edwards, 1995)(Fig. 3A).

The network was also characterized by a group of orthologs connecting the main clusters A and B. Three of these are involved in nitrogen (*nirA*) (Merrick and Edwards, 1995) (Fig. 3A), sulfur (*cysD*) (Pinto *et al.*, 2004) and central carbon metabolism (scoB) (Corthésy-Theulaz *et al.*, 1997) (Fig. 1B).

Globally, network analysis revealed that transcriptional activity of the community is structured around a selection of hub genes involved in central steps of nitrogen (*nirA*, *gudB*) (Fig. 3A) and sulfate assimilation (*cysD*, *cysN/C*), phosphorus acquisition (*ugpB*) (Fig. 3B) and carbohydrate metabolism (DLD, *scoB*), rather than around genes related to environmental stress resistance and damage repair (e.g. chaperones, proteases, SOD, *uvr*, *rec*), despite these genes being consistently active (Fig. 1).

## Conclusions

It is widely accepted that the extreme conditions in hot desert open soils limit both microbial and plant life (Pointing and Belnap, 2012; Makhalanyane *et al.*, 2015), and that microbial activity is spatially fragmented, temporally limited and water-driven (Pointing and Belnap, 2012; Belnap *et al.*, 2005; Collins *et al.*, 2014). We here show that there is a diverse and consistently active edaphic microbial community in open Namib Desert desiccated soils, dominated by non-photosynthetic bacteria, with a substantial actinobacterial and rhizobial component. Transcripts from all central metabolic pathway genes were detected, suggesting consistent metabolic activity during the study period. We therefore suggest that resistant microbial subpopulations remain active throughout long dry periods, rather than surviving in inactive states. Despite the observation of regular environmental fluctuations, no major diel changes were observed in prokaryotic activity, although a significant activation of fungal genes was noted during the night hours.

We identified the key microbial taxa responsible for the variety of strategies for carbon, nitrogen and phosphorus acquisition in the Namib Desert soil. Namely *Nitrospiraceae* (nitrate reduction), *Actinobacteria* (nitrite reduction, carbon fixation) and *alpha-Proteobacteria* (glycerol 3-phosphate assimilation) appeared as key functional members of the soil community.

Transcriptional network analysis of the community revealed a group of genes, involved in carbon, nitrogen, phosphorus and sulfur metabolism, in hub positions. This result supports our contention that the main driver of community functionality in Namib gravel plain soil under hyperarid conditions is nutrient assimilation rather than either environmental changes linked to day-night fluctuations (temperature, light, humidity) or the activity of stress resistance and repair genes.

Chemoautotrophic carbon fixation genes were among the most transcribed overall, indicating that this constitutes an important form of carbon assimilation under conditions where photosynthesis is restricted. We hypothesize that non-photosynthetic carbon fixation could be a strong adaptive factor in hyperarid, carbon-poor soils. Biochemical evidence is required to quantify the contribution of chemoautotrophy to the microbial community carbon flux.

Most notably, microbial N acquisition appeared to be limited to dissimilatory nitrate reduction. While some of the most transcriptionally active actinobacterial and rhizobial taxa are known to possess the capacity for dinitrogen fixation, there was no evidence that this was a significant contribution to nitrogen input budgets under desiccated soil conditions. We suggest that dinitrogen fixation processes may only become significant in times of higher water activity (such as after rainfall).

## Materials and Methods

### Sampling procedure

The sampling site was located in the gravel plains of the central Namib Desert (23°33′34′′S 15°02′25′′E), Namibia, approximately 56 km from the coast. The mean annual precipitation at the site is estimated at 25 mm, principally derived from nocturnal marine fog (Eckardt *et al.*, 2013). A 10 × 10 m experimental plot was sub-divided into 64 quadrats (Supplementary Fig. S1). Surface soils (0-4 cm) were collected at 6 hourly intervals (6:00, 12:00, 18:00, and 24:00 h) over three days from the 12^th^ to the 14^th^ April 2016. Two Hygrochron iButton sensors (Embedded Data Systems, Lawrenceburg KY, USA) were positioned at the corners of the plot, at ~2 cm depth, recording temperature and relative humidity at 4 minute intervals for the length of the experiment (Fig. S1). Soil respiration measurements were performed at the designated sampling times at four points within the plot using a LI-8100 IRGA (LI-COR Biosciences, Lincoln NE, USA), covering an area of 83.7 cm^2^ with a 3 litre chamber for 30 seconds (Supplementary Fig. S1). Photosynthetically active radiation (PAR) was measured using a photometric sensor (Quantum, LI-COR) at the same internal plot locations. Surface soil samples (0-4 cm) were collected at 6 hourly intervals (6:00, 12:00, 18:00, and 24:00 h) over three days from the 12^th^ to the 14^th^ April 2016. Three randomly selected quadrats were sampled at each time point (Supplementary Fig. S1). 20 g soil samples were immediately preserved on-site in RNAlater solution (Sigma-Aldrich, St. Louis MO, USA), temporarily stored at −20ºC at the Gobabeb Research and Training Center and during transport to the laboratory, and subsequently at −80ºC prior to total RNA extraction. An additional 400 g of soil for physicochemical analysis was collected in WhirlPak bags (Nasco, Fort Atkinson WI, USA) and preserved at 4ºC before physicochemical analyses. Soil pH, conductivity, cation exchange capacity (CEC), total nitrogen (%N), phosphorus (P), sodium (Na), potassium (K), calcium (Ca), magnesium (Mg), Chloride (Cl), Sulphate (SO_4_), ammonium (NH_4_) and nitrate (NO_3_) contents were analyzed by Bemlab (Pty) Ltd. (http://www.bemlab.co.za/; Strand, Western Cape, South Africa) using standard protocols.

### Total RNA purification

Soils from twelve physicochemically representative quadrats representing all sampling times were selected for RNA extraction (Supplementary Fig. S1, Table S2). Frozen, RNAlater-preserved soils were thawed at 4ºC, centrifuged at 14,500 rpm for 5 minutes and supernatants were discarded. 5 volumes of ice-cold 10 mM Tris-HCl 1 mM EDTA pH 6.5 buffer containing 100 mM NaH_2_PO_4_ were added to the soil to remove RNAlater salts. The supernatant was discarded after rapid (4 min) centrifugation at 4ºC. 0.5 volumes lysis buffer (5% CTAB, 0.7 M NaCl, 240 mM KH_2_PO_4_, pH 8) and an equal volume of TRI Reagent (Sigma-Aldrich) were added, and samples were vortexed at high speed for 30 seconds. RNA purification proceeded according to the manufacturer’s instructions. Extracted and purified total RNA was incubated with DNAseI (Invitrogen, Carlsbad, USA) and precipitated in the presence of 20% isopropanol and 15 ng glycogen co-precipitant (GlycoBlue, Invitrogen). RNA concentration and integrity were analysed using a Nanodrop 2000 spectrophotometer (Thermo Scientific, Waltham, USA) and 1% agarose gel electrophoresis. The absence of RT-PCR inhibitors was tested using the Transcriptor cDNA Synthesis Kit v9 (Roche, Indianapolis IN, USA) and universal bacterial 16S rRNA gene primers E9F (5′-GAGTTTGATCCTGGCTCAG-3′) and U1510R (5′-GGTTACCTTGTTACGACTT-3′) (Reysenbach and Pace, 1995; Hansen *et al.*, 1998).

### Library construction and sequencing

1 μg DNA-free total RNA from each sample was used for each sequencing library. Due to low RNA yields, we combined RNA from two Day 2 6:00h quadrats (Supplementary Table S1). Construction of rRNA-depleted libraries was carried out with the ScriptSeq Complete Gold Kit (Epidemiology) (Epicentre, Madison WI, USA), following the manufacturer’s instructions. Briefly, rRNA was removed by hybridization with bead-immobilized prokaryotic and eukaryotic 28S, 23S, 18S, 16S, 5.8S, 5S, mt16S and mt12S probes prior to RNA fragmentation and reverse transcription with tagged random hexamer primers. cDNA was amplified with TruSeq adaptors containing unique indexes (ScriptSeq Primer Set 1, Epicentre) for 15 PCR cycles. Libraries were purified using AMPure XP beads (Beckman-Coulter, Brea, USA) and final yields were measured with the High Sensitivity dsDNA reagents on a Qubit 2.0 fluorometer (Invitrogen). Multiplexed samples were quality and size analyzed in a High Sensitivity D1000 TapeStation (Agilent, Waldbronn, Germany). Libraries were single-end sequenced in a NextSeq500 v2 platform using the NextSeq 500/550 High Output v2 kit (Illumina, San Diego, USA). RNA-seq data were deposited in the ArrayExpress database (www.ebi.ac.uk/arrayexpress) and can be accessed using the reference E-MTAB-6601.

Read quality trimming was performed using Prinseq-lite v0.20.4 (Schmieder and Edwards, 2011) on both read ends with a mean Phred value of ≥30 in a 6 base sliding window. Reads shorter than 40 bases after trimming were discarded. rRNA and human-derived reads were removed from the dataset using Bowtie2 (Langmead and Salzberg, 2012) with a database of large- and small ribosomal subunit genes from SILVA (https://www.arb-silva.de/), 5S rRNA genes from the 5SRNAdb repository (Szymanski *et al.*, 2016) and the GRCh38 human genome primary assembly (ftp://ftp.ncbi.nlm.nih.gov/genomes/archive/old_genbank/Eukaryotes/vertebrates_mammals/Homo_sapiens/GRCh38/seqs_for_alignment_pipelines/GCA_000001405.15_GRCh38_no_alt_analysis_set.fna.bowtie_index.tar.gz).

### Analysis of sequencing reads

Functional and taxonomic profiling, and differential transcription analyses were performed using R version 3.3.3 (R Core Team, 2017). Read taxonomy was inferred from the NCBI Referece Sequence (RefSeq) database, and function was assigned based on the Kyoto Encyclopedia of Genes and Genomes (KEGG) Orthologs (KO) database (Kanehisa *et al.*, 2016) using the MG-RAST server (http://metagenomics.anl.gov/) (Meyer *et al.*, 2008). Read count tables were assembled and analysed for temporal expression changes using the EdgeR package (Robinson *et al.*, 2010) for all genes with > 1 count per million (cpm) in at least 3 libraries (n=11). Normalized KEGG ortholog counts were fitted to a generalized log-linear model (glmQLFit function) (Robinson and Oshlack, 2010; McCarthy *et al.*, 2012; Lun *et al.*, 2016), and pairwise comparisons between all time points were performed. Additionally, grouped “day” samples from 12:00 and 18:00 h were compared to “night” samples from 24:00 and 6:00 h. KOs were considered significantly differentially expressed between time points below a false discovery rate (FDR) corrected p-value threshold of 0.05.

Orthologs with average log_2_CPM values > 7 were used to construct a transcriptional network (Fig. 5), excluding KEGG categories Human Diseases and Organismal Systems. This threshold was selected as being below the common dispersion value calculated during differential expression analysis in order to reduce interference from high-variance, low-abundance transcripts. A transcriptional network was constructed using MENA’s (Deng *et al.*, 2012) RMT-based modeling with a correlation cutoff of 0.900 (p ≤ 0.005). Co-transcription was determined using Pearson’s correlation coefficients across libraries (n=11). The network was visualized using Cytoscape v. 3.5.1 (Shannon *et al.*, 2003).

## Declarations

### Data availability

The dataset supporting the conclusions of this article is available in the ArrayExpress repository, https://www.ebi.ac.uk/arrayexpress/ (accession no. E-MTAB-6601).

### Author contributions

C. L.-S., J.-B. R. and D. A. C. conceived the experiment. G. M.-K. provided logistical support and field advice in the Namib Desert. C. L.-S. performed all experimental work and bioinformatic analysis of the sequencing output. C. L.-S., J.-B. R. and D. A. C. participated in the interpretation of results and writing of the manuscript. D. A. C. provided funding.

## Acknowledgements

The authors wish to thank the Gobabeb Research and Training Station personnel for their assistance, advice and for providing access to their facilities during the sampling process.

